# Natural variability and individuality of walking behavior in *Drosophila*

**DOI:** 10.1101/2023.11.14.567019

**Authors:** Vincent Godesberg, Till Bockemühl, Ansgar Büschges

## Abstract

Insects use walking behavior in a large number of contexts, such as exploration, foraging, escape and pursuit, or migration. A lot is known on how nervous systems produce this behavior in general and also how certain parameters vary with regard to walking direction or speed, for instance. An aspect that has not received much attention is if and how walking behavior varies across individuals of a particular species. To address this, we created a large corpus of kinematic walking data of many individuals of the fruit fly *Drosophila*. We only selected instances of straight walking in a narrow range of walking speeds to minimize the influence of these high-level parameters, aiming to uncover more subtle aspects of variability. Using high-speed videography and automated annotation we captured the positions of the six leg tips for thousands of steps and used principal components analysis to characterize the postural space individuals used during walking. Our analysis shows that the largest part of walking kinematics can be described by five principal components (PCs). Separation of these five PCs into a 2-dimensional and a 3-dimensional subspace was found to divide the description of walking behavior into invariant features shared across individuals and features that relate to the specifics of individuals; the latter features can be regarded as idiosyncrasies. We also demonstrate that this approach can detect the effects of experimental interventions in an unbiased manner and that general aspects of individuality, such as the average width of the individual walking posture, can be described.

**Summary statement:** The kinematics of walking behavior in *Drosophila* can be decomposed into general aspects of motor output and idiosyncrasies of individual flies.

## Introduction

Legged locomotion, more commonly subsumed under the term walking, is found in most terrestrial animal groups. Walking is used in a diverse set of behavioral contexts, such as exploration and foraging, escape, pursuit, mating, and migration, making it a central component of an animal’s behavioral repertoire. The diversity of these contexts also means, however, that walking has to be highly adaptable and flexible. Consequently, the task- and situation specific neuronal control of walking behavior plays an important role for its proper execution.

The general and specific neuronal control and the kinematics of walking behavior have been extensively studied in a large variety of arthropods (Nirody, 2021; Nirody, 2023), particularly in insects, ranging from small insect species, like fruit flies (*Drosophila melanogaster,* DeAngelis et al., 2019; Mendes et al., 2013; Mendes et al., 2014; Strauß and Heisenberg, 1990; Wosnitza et al., 2012) or desert ants (*Cataglyphis,* Pfeffer et al., 2019; Wahl et al., 2015; Zollikofer, 1994), to large ones, like cockroaches (*Periplaneta americana,* Couzin-Fuchs et al., 2015; Delcomyn, 1971; Delcomyn, 1989), locusts (*Schistocerca gregaria,* Burns, 1973; Niven et al., 2010; Pearson and Franklin, 1984), or stick insects (*Carausius morosus,* Cruse, 1976; Dallmann et al., 2016; Du□rr and Ebeling, 2005; Gruhn et al., 2009). There is a large body of knowledge in these groups, ranging from the sensorimotor control of individual legs, to how functional interleg coordination of the six legs is achieved, to high-level descending and central neuronal control (Bidaye et al., 2018; Cruse, 1990; Dürr et al., 2004). To date, a number of common features governing the neuronal and kinematic aspects of insect walking have been identified. Walking speed, as a major aspect, seems to be generally controlled by changes in stance duration, while stance amplitude and swing duration are largely kept constant (DeAngelis et al., 2019; Wosnitza et al., 2012). These changes in stance duration are, in turn, accompanied by systematic changes in interleg coordination. Unlike larger vertebrates, however, which transition between distinct gaits in a speed-dependent manner (Diedrich and Warren, 1995; Hoyt and Taylor, 1981), insects exhibit a smooth and monotonic continuum of interleg coordination patterns (Szczecinski et al., 2018; Wosnitza et al., 2012). These systematic speed effects and other invariant features are generally present in walking insects as an evolved phenotypical trait; however, the neuronal control and kinematics of walking must have exhibited hereditary inter-individual and inter-species variability over evolutionary times, thereby adapting to co-evolving traits like body size and weight, leg morphology, and ecological demands. Indeed, a previous study on the evolution of walking behavior in a large set of drosophilids showed that there exist systematic differences in walking behavior across species and strains. These differences evolve and diverge rapidly in closely-related species, but also re-converge to shared features in more distantly-related ones (York et al., 2022). How variable this behavior is within and between individuals of a given species or strain, however, is still largely unexplored. Filling this gap should provide a better understanding of the ranges and covariations of the fundamental parameters defining walking behavior, thereby enabling more precise investigation of the general principles underlying the motor control of insect walking.

Walking behavior differs in multiple parameters between individual flies, such as average posture, preferred coordination pattern, walking speeds, or degree of intraindividual variability, as anecdotal and qualitative observations from our previous studies indicated. However, a quantitative description of these differences is still lacking. To explore this aspect in greater detail, in the present study we wanted to explicitly acknowledge variability as an important aspect of walking behavior on the inter- and intra-individual level and to find a less biased and more comprehensive way of measuring, describing, and interpreting the observed behavioral variability. We investigated the natural intra- and interindividual variability of low-level kinematic parameters of walking behavior in the fruit fly *Drosophila melanogaster*. To control for and exclude known influences of walking speed and curve walking on kinematics, we focused on straight walking at intermediate speeds in a large set of male flies (n = 88), each of which spontaneously produced a large corpus of walking behavior in an unrestrained free-walking paradigm. Using high-speed video recording and largely automated annotation based on deep learning methods (DeepLabCut, DLC Mathis et al., 2018), we extracted the positions of two body markers, as well as the six tarsal tips from all video frames in these straight sequences, automatically determined positions and times of lift-off and touch-down events of the legs, and calculated walking speed and interleg coordination for all step cycles. In total, the data set we created in this way contained more than 36,000 steps for each leg across all individuals. We used principal components analysis (PCA) to find a compact description of this large data set and systematically explored correlations and the variability between the kinematics of legs on an individual basis as well as across individuals.

Application of PCA revealed that most of the kinematic variability in the data set (approx. 80%) is contained in the first five PCs, which will be elaborated upon in the following. We show that two subsets of these five PCs describe inter-individually applicable dynamics of interleg coordination on the one hand and individual characteristics of walking behavior, what might also be called idiosyncrasies, on the other. The first subset contains two PCs which mainly capture interleg coordination-specific aspects of walking – how tripod-like a particular movement pattern is or the general repetitive sequence of alternating swing and stance movements of individual legs. In contrast, the contribution of a second subset of three PCs relates more closely to how individuals differ from each other in the way they walk. The data for different individuals occupy different regions within this PC subspace, highlighting inter-individual differences described by these three PCs. At the same time, the same flies are indistinguishable in the subspace related to interleg coordination, supporting the notion that these two PCs describe universal aspects of intra-individual variability, such as interleg coordination. The importance and applicability of these two subsets of PCs is further substantiated by (1) relating them to a quantitative measure of tripod coordination strength (TCS), (2) a use case for quantifying and comparing changes in walking behavior induced by optogenetic inhibition of proprioceptive sensory structures in the legs, and, finally, (3) deriving a one-dimensional measure for postural width. Taken together our results suggest that the variability observed in straight walking flies is highly systematic and that PCA is a suitable approach for the quantification of and decomposition into idiosyncrasies, inter-leg coordination patterns, and effects of experimental interventions.

## Materials and Methods

### Fly strains and husbandry

Male flies of the wild-type strain Berlin-K (Bloomington Drosophila Stock Center (BDSC, #8522) were used for those experiments which formed the basis for PCA (see below). Inhibition experiments (see below) were performed with F1 flies resulting from crosses between *iav*-Gal4 (O’Dell and Burnet, 1988) (BDSC #52273) and UAS-GtACR1 (Govorunova et al., 2015; Mohammad et al., 2017) (BDSC #92983). Wild-type flies were raised on 12h/12h light/dark cycle, while transgenic flies were raised in the dark to prevent premature activation of GtACR1 channels and potential adaptation prior to experiments. All flies were kept at 25°C and approximately 60% humidity on a standard food medium (Backhaus et al., 1984). To improve the function of GtACR1, transgenic flies additionally had 60 µg of all-trans-retinal in their food for at least three days prior to the experiments. To control for age-related effects all animals used in this study were five days old. Prior to an experiment flies were isolated and starved for approximately 24 hours to increase their walking activity. To prevent desiccation, flies had access to moist tissue paper during this period of isolation.

### Experimental setup

The experimental setup described here is largely identical to the one used in a previous study (Chockley et al., 2022). However, for clarity we describe it in detail here again. The recording arena (Fig. 1A) consisted of an inverted glass petri dish (diameter: 60 mm) as the walking substrate and a watch glass (diameter: 100 mm) as the lid. This arrangement formed a closed chamber with a curved dome tapered towards the edge of the petri dish, similar to an inverted FlyBowl (Simon and Dickinson, 2010). The inner side of the watch glass was coated with SigmaCote (SL2, Sigma-Aldrich, St. Louis, Missouri, USA). This resulted in a hydrophobic surface on which flies found less grip; walking on the ceiling was thus reduced. The petri dish and the watch glass were placed in a custom-made plastic holder with a cutout that allowed for video recordings from below (holder not shown in Fig. 1A). To record flies walking on the glass substrate we used a camera (model: VC-2MC-M340, Vieworks, Anyang, Republic of South Korea) equipped with an object-space telecentric lens (focal length 55 mm, model: Computar TEC-55, CBC America, Cary, North Carolina, USA). The telecentric lens provided an orthographic projection and thus reduced image position-dependent changes in apparent fly posture. The camera was located on the side of the setup and its view was directed at the experiment chamber from below via a surface mirror tilted at an angle of 45°. To increase video resolution the camera view was focused on a square area in the center of the surface of the walking substrate (30 mm side length, resolution 1000 by 1000 pixels, 33.3 pixels mm^-1^ or 30 µm pixel^-1^). The typical body length (BL, defined as the distance between the anterior end of the head and the tip of the abdomen) of a fly of approximately 2 mm corresponded therefore to 65 - 70 pixels; this was sufficient to image all body parts annotated during analysis with sufficient accuracy, particularly the tarsal tips (Fig. 1B, left). The scene was illuminated with 60 infrared (IR) LEDs (wavelength: 890 nm, opening angle: 20°) arranged in a concentric ring around the chamber. IR light from these LEDs was emitted mainly parallel to the arena surface. Thus, from the perspective of the camera, only the fly reflected any appreciable amount of light, resulting in a strong contrast between it and the background (see Fig. 1B). Contrast was further enhanced by adding a visible-light filter (upper cut-off frequency 790 nm) to the camera’s lens, thus eliminating any ambient light from the room or optogenetic inhibition (see below). Video data was acquired at 200 Hz and a shutter time of 400 µs; this low shutter time prevented motion blur and ensured detectability of the leg tips even during very fast movements. Acquisition of individual video frames during this shutter time and IR illumination were synchronized with a pulse generator. For inhibition experiments (see below) we added a second LED ring around the chamber. This ring consisted of 60 green LEDs (wavelength: 525 nm) whose light was directed at the recording chamber. These LEDs could be switched on and off programmatically during an ongoing experiment via a multi-function I/O device (USB-6001, National Instruments Corporation, Austin, TX). The experimental setup was custom-designed and built (Electronics workshop, Department of Animal Physiology, University of Cologne).

**Figure 1:**
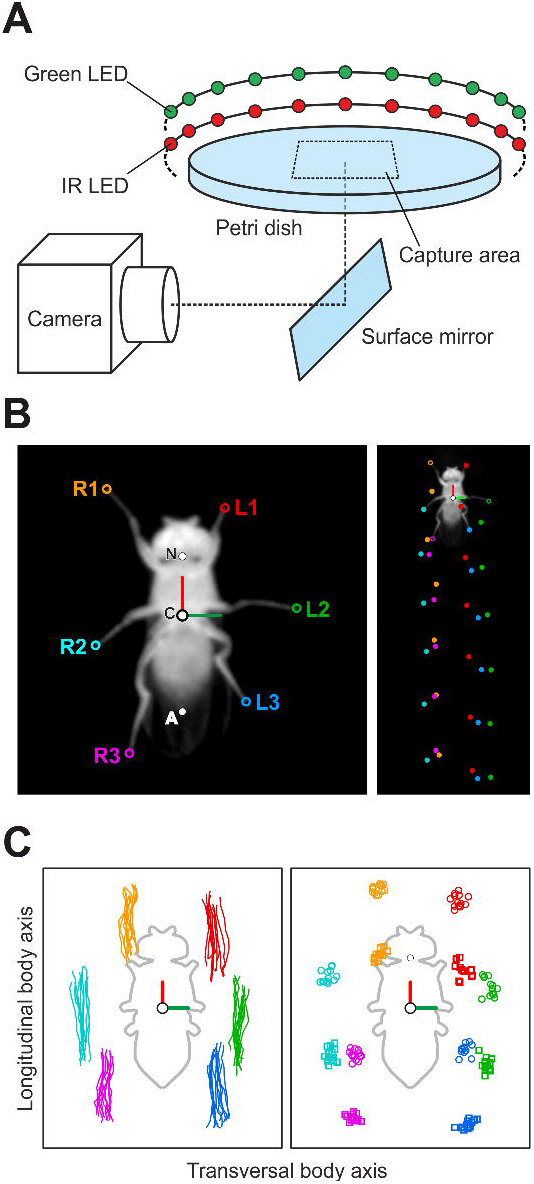
Setup and data acquisition. (A) Schematic of the experimental setup. The watch glass covering the petri dish is not depicted. (B) Examples of the body-centered bottom view (left) and the camera view (right) with annotated leg tips (right legs R1 - R3, and left legs L1 - L3), abdomen (AT), center of the body (C) and neck (N). (C) Examples of stance trajectories (left) and tarsus extreme positions (right) of one walking bout after detection of lift-off and touch-down positions. Round markers: AEPs, square markers: PEPs. Fly silhouette (gray outline) for reference.

### Behavioral experiments

Prior to an experiment, single flies were aspirated into a tube and then transferred into the arena; no CO_2_ was used for this step. After the transfer, flies were allowed to acclimate to the setup for 5 minutes prior to the start of video acquisition. The total duration of an experiment was up to 3 hours. During this time, flies walked spontaneously in the chamber and frequently crossed the capture area. For the complete duration of an experiment, video data was acquired continuously and the last 1000 frames (equivalent to 5 seconds) were stored in a ring buffer. Custom-written software functions evaluated the recorded frames in real-time and determined if the fly was present in a particular video frame and if it had produced a continuous walking track with a minimum length of 3 BL and a minimum walking speed of 2 BL s^-1^. Once the fly had produced such a track and then stopped or left the capture area the contents of the frame buffer were committed to storage as a valid trial. After this, acquisition automatically started anew. Note, that at this time no additional selection criteria (such as curvature of the trajectory or particular walking speeds, for instance) were applied to determine whether a trial was valid or not. The set of videos acquired in this way merely served as a large and relatively unconstrained initial set of walking behavior (see next section on further criteria for data inclusion).

The inhibition experiments used for evaluation of the descriptive power of the principal components (see below) were performed in the same setup. For this purpose, the experimental paradigm was extended as follows. Flies in these experiments either walked in the dark (wild-type control condition) or under green-light illumination (inhibition condition). Trials in the dark and in the light were alternated; once the fly had produced a valid trial in the dark, the green-light illumination was switched on and the system was primed to record the next valid trial. If the fly did not produce a valid trial within the first 60 seconds of the green-light condition, the light was switched off and data acquisition was suspended for 30 seconds to prevent adaptation to the inhibition and potentially harmful effects of prolonged exposure to the green light. After this cooldown period, the green light was switched on again, and data acquisition was resumed normally. Once a valid trial had eventually been recorded in the inhibition condition, the light was switched off again and the process was repeated. Data from these inhibition experiments were then sorted into control trials (recorded in the dark) and inhibition trials (recorded during green-light illumination). Video acquisition, online data evaluation during experiments, and general high-level hardware control were implemented with custom-written software in MATLAB (2018b, The Mathworks, Natick, Massachusetts, USA).

### Processing of video data

Because curve walking has a strong influence on leg kinematics in walking insects (Du□rr and Ebeling, 2005; Gruhn et al., 2008; Jander, 1985) we restricted all further analyses to straight walking. Instances of this in the complete set of recorded walking trials (see previous section) were detected with custom-written algorithms, whose parameters were determined empirically (see Fig. S5) and only segments of video data associated with straight walking were extracted from valid trials for further analysis. Generally, in each video frame of these segments the tarsal tips of all six legs and two additional body parts (the neck and the posterior tip of the abdomen) were detected with DeepLabCut (DLC) (Mathis et al., 2018), for a total of eight different body parts (Fig. 1B). To accelerate data processing and improve robustness of DLC we first detected the general position of the fly within a video frame using a simple threshold operation, conversion to a binary image, and, finally, calculation of the centroid of the largest contiguous region of white pixels (which corresponded to the fly). Using this position, we then cropped the fly from the video. With these cropped and fly-centered views we used three different instances of DLC in a two-step analysis. The first step was to detect the neck and abdomen of the fly. These positions were used to define a fly-centric coordinate system (Fig. 1C); all data presented in this study (apart from the identification of swing and stance phases) are based on these body-centric coordinates. The positions of all body parts were then normalized to each fly’s body length to allow for body size-independent comparisons between flies. We also used the neck and abdomen positions to rotate the cropped views and align the fly’s longitudinal axis vertically. These cropped and rotated data were then used in the second DLC analysis, in which the tarsal tips of each body side were detected by two independently-trained instances of DLC, one for the left and one for the right legs. In general, DLC performance was very good; to ensure highest accuracy, however, we also visually inspected all automatically generated annotations for errors and corrected these manually, where necessary.

Swing and stance phases of all legs were determined automatically based on the respective speeds at which tarsal tips moved in an arena-centric coordinate system: whenever a tarsus is stationary in this coordinate system, i.e. it co-moves with the ground, we assumed the leg to be in stance phase (Fig. 1B, right). Conversely, movements of more than 1.5 pixels per frame were empirically defined as swing phase activity (see Fig. S3). A transition between stance and swing phase was defined as lift-off event. The last position of the tarsal tip on the ground before lift-off was defined as the posterior extreme position (PEP) for that step (Fig. 1C, right). Conversely, a transition between swing and stance phase was defined as touch-down event; the first tarsal position with ground contact associated with this event was identified as the anterior extreme position (AEP) (Fig. 1C, right). The time of onset of a particular step was defined as its lift-off, and a complete step of a leg was defined as its movement between two consecutive lift-off events, i.e. a swing phase followed by a stance phase. In contrast to steps of individual legs, step cycles (SCs) were defined as follows: start and end of an SC were determined by the respective step of the right middle leg, which was selected arbitrarily for this purpose. All six tarsal tip positions for this interval comprised the data of one SC. Since the stepping period of the six legs in straight walking flies is almost identical and constant for small time windows, each leg completes its own cycle during an SC, although they all start and end at individual positions and phases, respectively. In other words, all six leg tip positions at the beginning and the end of an SC are usually highly similar, not just for the reference leg.

The walking speed associated with a step or an SC was defined as the average walking speed of the animal between on- and offset and was used to allow for the selection of steps and SCs within a certain range of walking speeds for analysis. Data used in the present study was based on steps and SCs whose associated walking speed was between 5 and 7 BL s^-1^ (see Fig. S1 for all speed ranges of all individuals). We restricted the range of walking speeds in this way to facilitate comparability between individuals. Furthermore, previous studies have shown that walking speed has a strong and systematic influence on many of the kinematic parameters investigated here (Mendes et al., 2013; Strauß and Heisenberg, 1990; Wosnitza et al., 2012); this general influence might have an unwanted effect on the analysis if individuals walk at different preferred speeds.

### Principal component analysis (PCA)

PCA is a tool for dimensionality reduction and can be used to find linear correlations of multiple parameters in high-dimensional data sets. Mathematically, calculating PCA is identical to finding the eigenvectors and eigenvalues of a data set’s covariance matrix (Manly and Alberto, 2019). The principal components (PCs) form a new coordinate system that is aligned with the directions of highest variability in the data set. Importantly, PCs are ordered according to their relevance, i.e. the relative amount of variance they describe.

Here, we applied PCA to the positions of all six leg tips during SCs for 88 individual flies. 30 SCs between 5 and 7 BL s^-1^ for each of these 88 flies were used as the basis for PCA. Resampling and interpolation were used to acquire exactly 100 postures for each SC in the analysis, resulting in matrices of 3000-by-12 data points per fly, with each row representing x and y-coordinates of the six leg tips and a total of 30 (SCs) times 100 (normalized number of data points per cycle) rows. The final data matrix for PCA comprising data from all flies contained 2,640 SCs, represented by 264,000 data points with 12 parameters each. Prior to the analysis, each column (equivalent to one dimension or parameter) was standardized to a mean of 0 and unit variance (equivalent to the calculation of z-scores). PCA was carried out on this standardized matrix in MATLAB 2018b (function *pca.m*).

Here, PCs describe spatial covariations of the positions of all six tarsal tips and can readily be interpreted as shifts in tarsal positions when multiplied by a non-zero factor. For this, individual PCs were visualized by varying their values systematically from minus 2 to plus 2 of their respective standard deviations before transferring the data back into the original parameter space. The resulting positional changes are depicted as arrows to indicate the direction of covariation. In mathematical terms, these arrows correspond to the *loadings* of each PC. For subsequent data analysis, the original data was transformed either in its entirety (264,000-by-12 matrix) or on a per-fly basis (3000-by-12) into the new coordinate system established by the PCs. In the context of PCA, these transformed data are typically also referred to as *scores*.

### PCs and tripod coordination strength (TCS)

The fraction of variability described by a PC for one complete cycle of both tripod groups was compared to the respective coordination pattern. Tripod coordination strength (TCS, as used in (Ramdya et al., 2017; Wahl et al., 2015; Wosnitza et al., 2012)) was used to quantify the synchronicity of the swing activity in the two tripod groups. Briefly, the TCS was calculated as follows: for each tripod group (a set of ipsilateral front and hind legs and the contralateral middle leg), the time in which all three legs were simultaneously in swing phase was divided by the time from the earliest swing onset to the latest swing termination in any of these three legs. Hence, a perfect overlap of all three swing phases resulted in a maximal TCS value of 1 and would correspond to canonical tripod coordination. The minimal TCS value of 0 was assigned in cases of no overlap between swing phases in a tripod group. The TCS values of the tripod groups were averaged and the fractions of variability described by each principal component for all positions during the movements of these two tripod groups were calculated. This was achieved by transferring all leg tip positions in this respective time frame into the principal component space and measuring the fraction of variability described by each PC.

### Evaluation experiments

To test the suitability of the PCA based approach for the description and analysis of idiosyncrasies in *Drosophila* walking behavior we used an optogenetic approach. For this purpose, the Gal4-UAS-system was used to express GtACR1, an anion-selective channelrhodopsin, in a group of mechanosensory neurons in the legs. When activated optogenetically, GtACR1 inhibits neurons in which it is expressed (Govorunova et al., 2015; Mohammad et al., 2017). We used the transgenic *iav*-Gal4 line (O’Dell and Burnet, 1988) to target all chordotonal organs, including the femoral chordotonal organ (fCO), the largest sensory organ in the fly’s legs. The resulting transgenic flies have been shown to exhibit a systematic and noticeable phenotype in walking behavior in an inhibition paradigm (Chockley et al., 2022). We used this to evaluate whether we can detect these known effects in the PCA approach explored here. For this, a minimum of 30 SCs each for dark (control) and light condition (inhibition) for individual flies was compared regarding the mean tarsal trajectories and the shift observed in the respective mean positions in PCs 2, 4, and 5. We tested the observed effect size for significance by comparing it to a bootstrap analysis performed on the original wild type data set: for 14 individual flies we randomly selected 2 unique sets of 30 SCs each and compared these two sets with each other as we did compare the two conditions (control and inhibition) for the transgenic flies.

### Symmetry axis in subspace of PCs 2, 4, and 5

The evidence that PCs 2, 4, and 5 describe inter-individual differences in leg kinematics and posture (see Results section) suggests that these PCs describe more general differences in the mean posture of individual flies. Among other aspects, this might refer to how sprawled the posture of an animal is or individual-specific distances between AEPs and PEPs, for instance. To give a comprehensible example, we systematically searched for an axis in the subspace of PCs 2, 4, and 5 along which the posture changes symmetrically with respect to the left and right body side. This axis was supposed to go through the center of the three-dimensional subspace and respective postures were calculated from minus to plus five times the standard deviation. Symmetry was measured by calculating the RMSE between contralateral leg-pairs after projecting the positions of one body side to the other. The axis with the highest symmetry scores and the respective postures are shown in Figure 7.

## Results

We recorded 103 male flies of the wild-type strain Berlin-K. They performed a total of 36,942 straight walking steps with a median of 242 steps per fly with 10% and 90% percentiles of 93.8 and 841 steps, respectively (min: 28; max: 1717). 88 flies yielded 30 or more steps within the speed range of 5 to 7 BL s^-1^ targeted in our analysis and thus were included in the PCA. Restricting our analysis to this fairly narrow speed range was important, because many basic parameters of walking behavior, such as duty cycles, stepping frequency, step amplitude, phase relations, and even variability itself change systematically with walking speed. During the evaluation experiments, we recorded 28 flies resulting from crosses between *iav*-Gal4 and UAS-GtACR1. They performed a total of 9,694 straight walking steps in the control and inhibition condition with a median of 279.5 steps per fly with 10% and 90% percentiles of 108.9 and 726.6 steps, respectively (min: 43; max: 918). 14 of these flies produced 30 or more steps within the speed range of 5 - 7 BL s^-1^ for both conditions and thus were included in the analysis.

### Qualitative inspection shows flies walk in an idiosyncratic manner

By recording leg tip positions of unrestrained, straight walking flies we analyzed the inter- and intra-individual variability of walking behavior under these conditions. Despite being all male, of identical age, from the same isogenic stock culture, and tested under identical conditions, the flies analyzed here showed clear idiosyncrasies that can already be seen during qualitative visual inspection. To illustrate this, Figure 2 shows AEPs and PEPs (Fig. 2Ai - Aiii) as well as leg tip trajectories (Fig. 2Bi - Biii) of three exemplary individuals over the course of walking bouts lasting seven to eight step cycles. While generally similar, several distinguishing characteristics of tarsal tip kinematics can be identified, such as (1) the overall width of the complete posture or leg pairs (compare Fig. 2Bi and Biii, for instance), (2) the straightness of tarsal trajectories and their specific shapes, or (3) overall left-right symmetry. Qualitatively, it is also evident that intra-individual step-to-step (i.e. distance between AEPs and PEPs or leg tip trajectories in one leg) variability is lower as compared to inter-individual variability (i.e. difference in posture or asymmetries between Fig. 2Bi and Biii). These observations show that flies indeed have distinct idiosyncrasies in their walking behavior.

**Figure 2:**
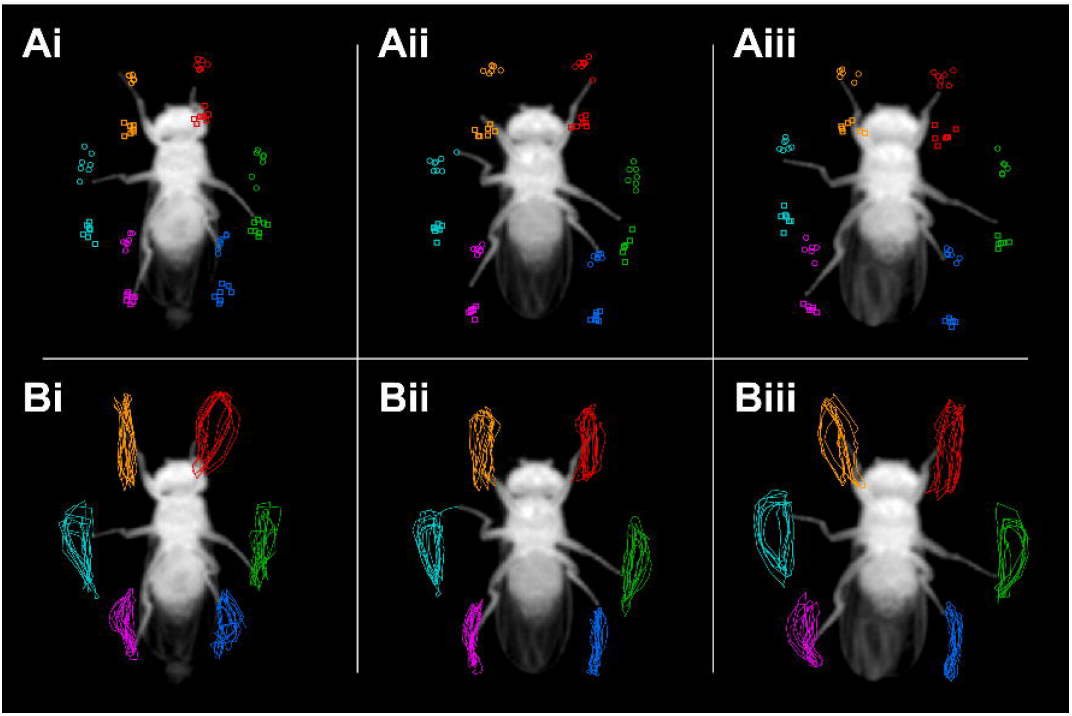
Qualitative inspection shows flies walk in an idiosyncratic manner. Here, inter-individual variability of leg kinematics in three exemplary individuals is shown (sub-indices i to iii refer to three different animals, respectively). (A) Anterior and posterior extreme positions (AEPs and PEPs) of single walking bouts. Round markers: AEPs, square markers: PEPs. (B) Complete tarsal tip trajectories of the same walking bouts as in A. Leg color coding: red = L1; green = L2; blue = L3; orange = R1; teal = R2; magenta = R3. These three flies are also referenced as exemplary flies in Fig. 5 and Fig. 7.

### PCA detects significant correlations of leg tip positions

We used PCA to test for and describe linear covariations of leg tip movements in a fly-centered coordinate system. Such covariations detected by PCA have either a systematic biological basis or are just random fluctuations. This can be tested by comparing the results with those of an additional PCA calculated for the same data set that was permuted randomly along the columns. The permutation retains all statistical features (e.g. mean, variance, or percentiles) of the original data set, but removes any covariations by reshuffling formerly correlated variables. The randomly permuted version of the original data set therefore is the perfect reference point for the strength of correlations which should be expected just by chance. Since our data set is fairly large, the fraction of variability described by each PC for the randomly permuted version approximated the reciprocal number of dimensions of 8.33% with a slight decrease from the first (8.42%) to the last PC (8.24%). Fractions of variability described by PCs 1 to 5 for the original unpermuted data set exceeded their respective reference value, hence they accounted for more variability than expected for uncorrelated data (Fig. 3A). In sum, these five PCs described 78.02% of the variability in the data set. PCs 6 to 12 might still describe non-random correlations, but increasingly unfavorable signal to noise ratios and the diminishing relevance of PCs with successively lower fractions of described variability made them less likely to capture important correlations. For all further analyses we therefore focused on these first five PCs.

**Figure 3:**
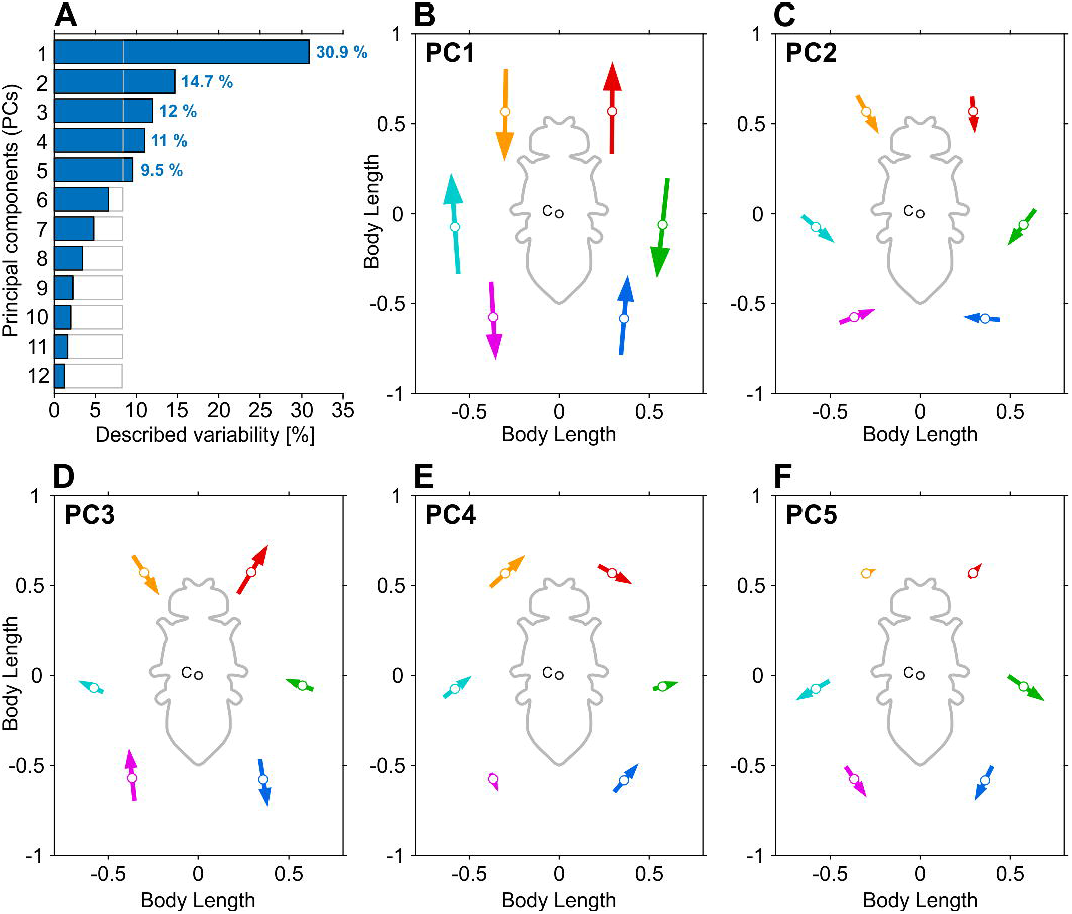
PCA finds significant correlations of leg tip positions. (A) Fractions of variability in leg tip kinematics described by each principal component. Filled blue bars: results based on PCA on the data set acquired in this study (number of flies: 88, number of steps per fly: 2,640). Gray open bars: results for the same, but randomly permuted data set (see results section). PCs 1 to 5 exceed the described variability expected based on the permuted data set (respective percentages shown at the bars). (B) to (F) Visualization of covariation of tarsal tip kinematics captured by PCs 1 to 5, respectively. White dots on arrows depict the mean positions of all data points in the analysis. Arrows indicate the directions and magnitude of covariations for a particular PC. C: center of the body. Leg color coding: red = L1; green = L2; blue = L3; orange = R1; teal = R2; magenta = R3.

Here, each PC intuitively describes how 2D positions of the tarsal tips covary. This covariation can either be a result of active leg movements during walking, i.e. protractions and retractions; a strong covariation in the two front legs, for instance, can be expected, because during anterograde movement of one front leg the other one generally moves posterograde. The second cause of covariation described by PCs is based on general postural difference between individuals; for instance, the degree of how sprawled an individual’s natural posture is will affect its six legs fairly evenly and will be detected as positional covariation that generally describes the distance of the tarsal tips from the body center. Other, less obvious, postural covariations are conceivable. Based on these considerations, individual PCs can be visualized as arrows to facilitate the comparison of the direction and magnitude of covariations (Fig. 3B - F). PC 1 (30.9% described variability, Fig. 3B) captured the counter-directed anterograde and posterograde movements of neighboring legs, either ipsilaterally or contralaterally, and therefore comprised the major component of forward locomotion. This PCs suggestively groups the six legs into two groups (R1, R3, and L2, as well as L1, L3, and R2, respectively) whose coupling and covariation corresponds to idealized tripod movement. However, the first PC described less than a third of the dynamics occurring in the leg tip movements. PC 2 (14.7%) shows shifts that are mirror symmetric along the longitudinal body axis for all three leg pairs. Front legs covary in the direction of movement, middle leg covariation is tilted by approximately 45 degrees pointing on the tip of the abdomen, and hind legs covary in parallel with the x axis. The directions of these covariations indicate that PC 2 might describe interindividual differences in postural width as observed between fly 1 and 2 (Fig. 2 Ai and Aiii). PC 3 (12%) again showed anterograde and posterograde directions of covariation for the front and hind legs (Fig. 3D). Here, however, the sign of covariation is negative on both body sides as compared to PC 1, and the middle legs do not display a comparable covariation in the direction of movement. Interestingly, PC 3 allows for a dissociation from the strict tripod coordination as defined by PC 1 (but see next section for further elaboration). PC 4 (11%) only exhibits asymmetric covariations. PC 5 (9.5%) again shows body side symmetric shifts which are almost perpendicular to the directions described by PC 2 (Fig. 3C), indicating that PC 5 also describes interindividual differences in posture.

### TCS reveals the relevance of PCs 1 and 3 for interleg coordination

Using tripod coordination strength (TCS) as a measure for how tripod-like a stepping sequence is we tested for correlations between inter-leg coordination patterns and scores of individual PCs. Insects use a continuum of interleg coordination patterns in contrast to the distinct gates observed in many vertebrate species (Szczecinski et al., 2018; Wosnitza et al., 2012). The selection of a particular pattern is mainly dependent on walking speed. High walking speeds are strongly associated with the so-called tripod coordination, while slower walking speeds effect coordination patterns that deviate from canonical tripod coordination (Szczecinski et al., 2018). Canonical tripod coordination (which is rarely observed in an ideal form) thereby corresponds to anti-phasic activity of two tripod leg groups (sets of ipsilateral front and hind legs and the contralateral middle leg). In this context, tripod coordination strength (TCS) is a single value that describes how similar a particular coordination pattern is to ideal tripod coordination (for a more detailed description see (Wahl et al., 2015; Wosnitza et al., 2012)). Here, we used TCS to test whether covariations described by PCs were indeed correlated with certain coordination patterns, as suggested by the findings described above. For this purpose, we first calculated TCS for all instances in which complete leg cycles of a particular tripod group were available. Then, where possible, we averaged the TCS values of two subsequent tripod cycles, effectively extending the TCS definition from single tripod groups to all six legs. Leg tip positions within the respective interval were mapped into PC space and relative fractions of variability described by each PC for this subset of data were calculated. The fraction of variability described by PC 1 was found to be strongly positively correlated with the TCS (Fig. 4B). In contrast, the described variance of PCs 2, 4, and 5 did not show a clear correlation with the respective TCS values (Fig. 4C, E, and F). PC 3 was negatively correlated with the TCS values, but this was less pronounced compared to PC 1 (Fig. 4D). Taken together, these results suggest that PCs 1 and 3 describe a substantial part of the dynamic and coordination-related aspects of walking behavior.

**Figure 4:**
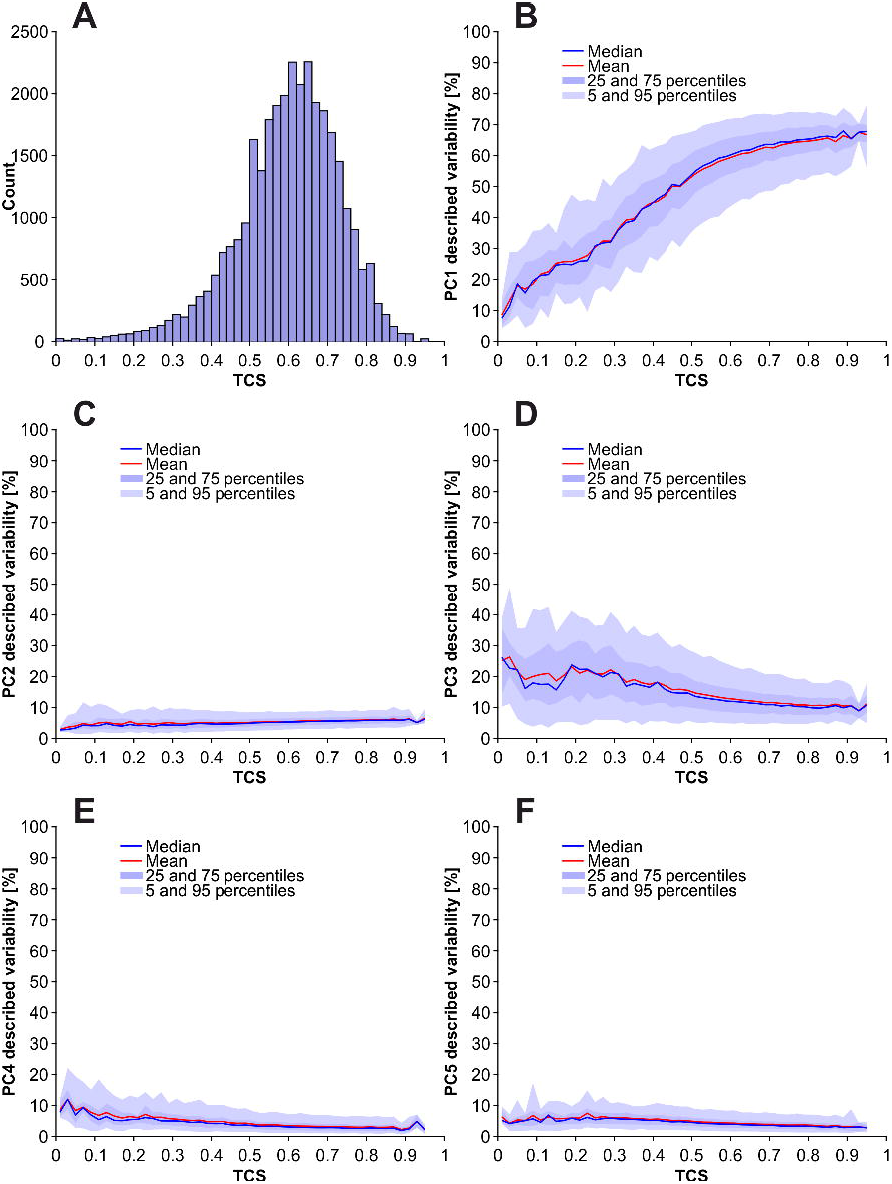
TCS reveals the relevance of PCs 1 (B) and 3 (D) for interleg coordination. (A) Histogram of TCS-values for all steps of all flies in the analysis pool. (B) to (F): TCS values plotted against described variability of PCs 1 to 5. TCS was calculated for consecutive cycles of both tripod groups. Scores for PCs 1 to 5 were determined by transferring the tarsal tip positions of respective steps into the PC space and calculating the relative described variability for each PC and each cycle. The higher the TCS the more tripod-like a particular cycle is.

### PCs 2, 4, and 5 describe inter-individual differences of leg kinematics and posture

Individual flies were compared in subspaces spanned by the first five PCs to check whether interindividual differences of fly walking behavior are resolved by these PCs (Fig. 5). One aspect not mentioned in the previous section is that fractions of variability described by a PC can be much smaller or larger for subsets of data from individual flies than for the complete set (Fig. 4B shows scores above 60%, Fig. 3A shows only 30.9% for PC 1 for the full data set). This can be explained by the fact that subsets of data from individual flies do not have the dimension of inter-individual variability, while variability resulting from regular forward and backward movements of leg tips, as well as different coordination patterns, are still present to a large degree. Hence, these intra-individual aspects make up for a larger fraction of variability in single-fly data, as shown for PCs 1 and 3 (Fig. 4B and D), compared to their scores for the complete data-set (Fig. 3A). Since this is another indication for the notion that PCs 1 and 3 mostly capture intra-individual aspects of variability, an open question is how inter-individual variability, in contrast, is resolved by PCs. To check for putative inter-individual differences in the PC space we plotted data from five exemplary individual flies in 2-dimensional PC subspaces (Fig. 5A - D). The respective subspaces spanned by pairs of PCs 2, 4, and 5 (Fig. 5A - C) show relatively clear separations between the flies, while the plot for PC 1 and 3 (Fig. 5D) displays an overlap of all flies in the analysis. Interestingly, data plotted in Figure 5 D forms an elliptic shape around the origin. We found that over the course of a step cycle the postural representation in this 2D subspace spanned by PCs 1 and 3 cycles the origin once (data not shown). The exact dimensions of the ellipse formed hereby is largely invariant for individuals, but strongly depends on the current interleg coordination pattern (data not shown). Boxplots in Figure 5E shows again that all 5 flies had individual signatures regarding their scores for PCs 2, 4, and 5, while scores for PC 1 and 3 were comparatively similar (median values for PC 1 and 3 were close to zero for all 5 flies). These results demonstrate that PCs 2, 4, and 5 describe some of the individual differences between flies, while the absence of separation in the subspace spanned by PCs 1 and 3 shows that this space describes features of the data that are largely invariant between individuals (such as general aspects of interleg coordination).

**Figure 5:**
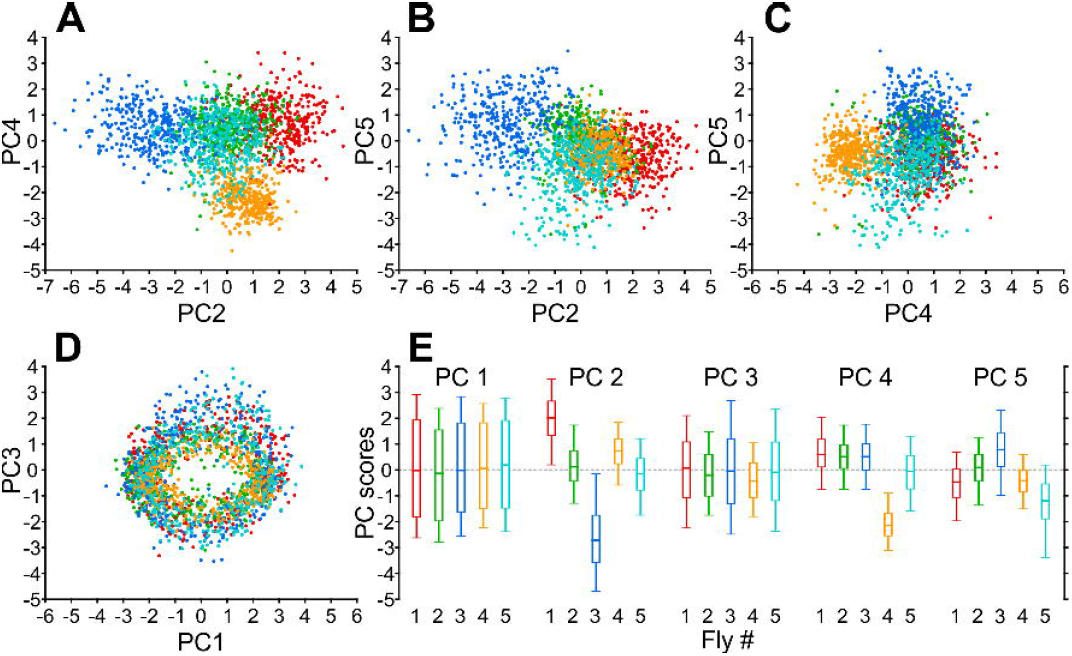
PCs 2, 4, and 5 describe inter-individual differences of leg kinematics and posture. (A) PCs 2 and 4, (B) PCs 2 and 5, (C) PCs 4 and 5, (D) PCs 1 and 3. Each dot represents a fly’s posture at a given time, selected randomly from the data which contributed to the analysis. Individual colors correspond to individual flies. (E) Boxplots indicating the distributions of scores of the five flies in A to D (same color code) for the first five PCs. Fly 1, 2, and 3 labeling refer to the data of exemplary flies presented in Fig. 2, the data for flies 4 and 5 have been added to further populate and exemplify the subspaces in A to D.

### An optogenetic inhibition experiment demonstrates the descriptive potential of PCs

An optogenetic inhibition experiment was performed to evaluate the potential of PCs 2, 4, and 5 to concisely describe systematic differences in posture and leg positioning. For this purpose, the *iav*-Gal4 driver line (BDSC #52273) was crossed with UAS-GtACR1 (BDSC #92983). The resulting F1 generation expressed the anion-selective channelrhodopsin GtACR1 in the neurons of chordotonal organs, including the femoral chordotonal organ (fCO), the largest sensory organ in insect legs. Previous experiments showed that inhibition of these sensory structures via optogenetic inhibition with GtACR1 leads to systematically and noticeably altered kinematics of walking behavior (Chockley et al., 2022), mainly elongated stance trajectories, but also some leg-specific changes. Here, we used these known and expected changes in kinematics to test if they can be detected and described compactly within the subspace spanned by PCs 2, 4, and 5.

For each fly with more than 30 steps in the control (dark) as well as the inhibition condition (light), the mean positions for both conditions were analyzed in the subspace spanned by PCs 2, 4, and 5 (Fig. 6A). The length of the vector describing the difference between dark and light condition was compared to a bootstrap analysis. For this bootstrap analysis, two sets of 30 steps each were randomly drawn from single flies of the original data set and plotted in the same way as the data from the inhibition experiments (Fig. 6B). The results for the inhibition experiments show no strong preference for shift directions for PCs 2 and 4, but a clear and consistent shift towards more positive values for PC 5 (Fig. 6Ai - Aiii). In contrast and expectedly, the bootstrap analysis resulted in more random shift directions whose magnitude was also smaller. The mean effect sizes measured for PCs 2, 4, and 5 (i.e. vector lengths) were plotted against the observed differences in the mean leg tip trajectories between control and inhibition, expressed as root mean squared error (RMSE), showing a positive correlation (Fig. 6Bi); the stronger the difference between control and inhibition on the level of actual leg tip kinematics, the larger the shift in PC space was. The difference in RMSE is mainly driven by the middle legs (green dots), matching the effect described by (Chockley et al., 2022). Figure 6Bii depicts the vector lengths for the inhibition experiments and the bootstrap analysis, showing that effect size was much larger on average for the inhibition experiments. PC 2 shows shifts to more negative values for 10 individuals, but also four in the opposite direction, although with much smaller amplitudes. A different effect on walking behavior more aligned with the covariations captured by PC 2 should result in a higher descriptive power. PC 4 shows even smaller and less consistent effects, likely because the inhibition of fCOs affected both body sides equally while PC 4 describes only asymmetric covariations (Fig. 3E). However, the clear and consistent shifts from dark to light condition in PC 5 demonstrate that PCs in general can be used to compare and quantify effects and their magnitude in a reduced number of dimensions.

**Figure 6:**
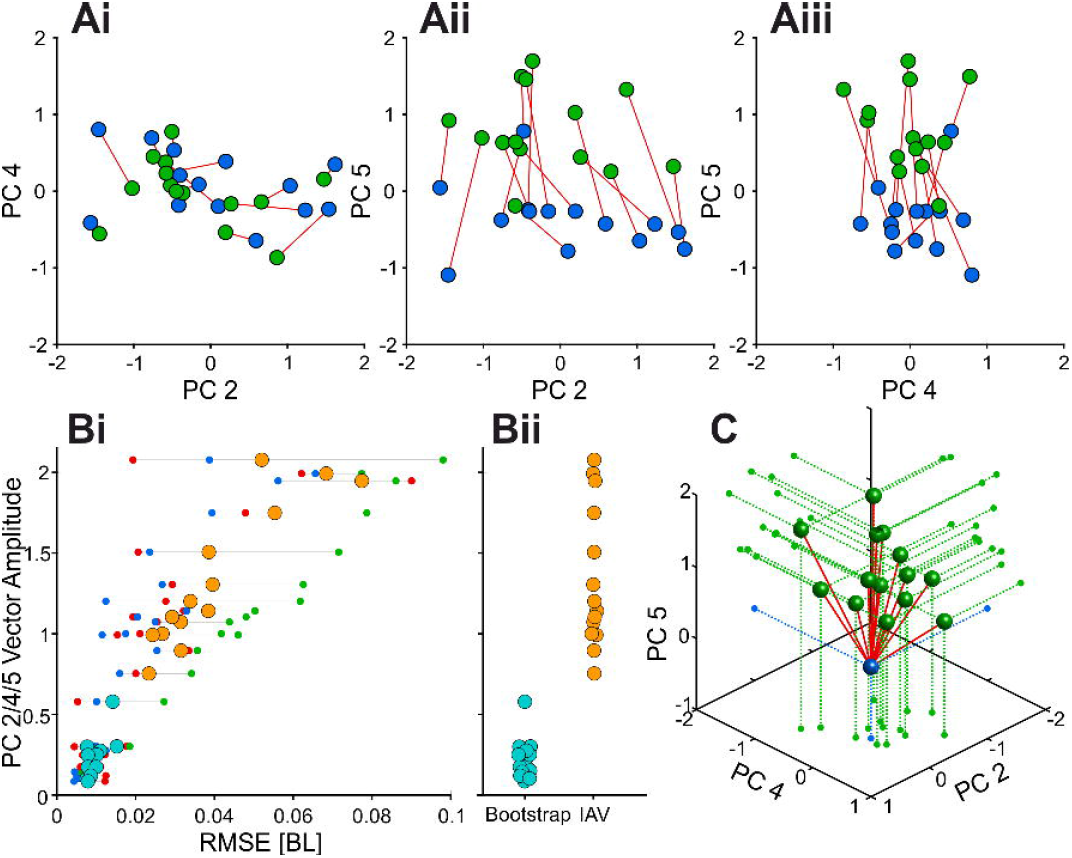
An optogenetic inhibition experiment demonstrates the descriptive potential of PCs. (A) 2-D representations of the inhibition effect in subspaces spanned by combinations of PCs 2, 4, and 5 (steps in dark: blue; steps in green light (fCO inhibited): green). Red lines connect mean positions of individual flies in the two conditions. (B) Vector amplitudes (i.e. effect size) of the inhibition experiments and the bootstrap analysis plotted against the RMSE of the leg tip trajectories. Teal dots depict results for the bootstrap analysis, orange dots for inhibition experiments. Red, green, and blue dots indicate the RMSE for front-, middle-, and hind legs. (C) 3-D representation of the inhibition effect (green) relative to the control condition (blue). For clarity, control conditions were set to the origin.

**Figure 7:**
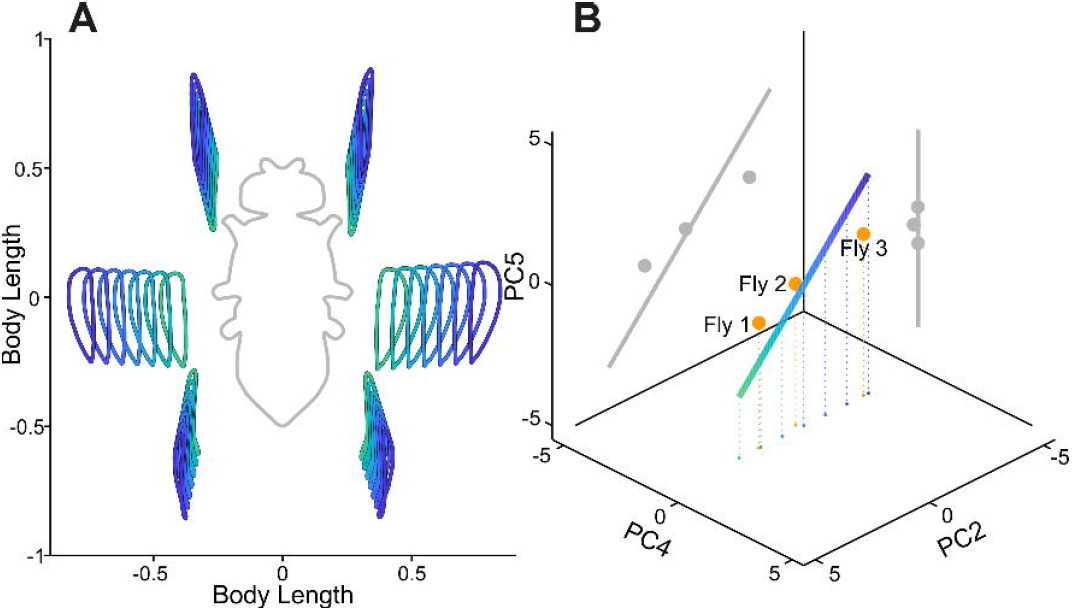
A symmetry axis demonstrates how individual postures are encoded in PCs 2, 4, and 5. (A) Leg tip trajectories for different points on the (B) symmetry axis in the subspace of PCs 2, 4, and 5 describing systematic differences in postural width shown in (A). Color code in A corresponds to color shading along the axis in B. Orange dots in B depict the mean positions of flies 1, 2, and 3 introduced in Fig. 2 and also referenced in Fig. 5. Gray: projections of the three-dimensional sequence of explored combinations into two-dimensional subspaces spanned by PCs 2 and 5 as well as PCs 4 and 5, respectively.

### A symmetry axis demonstrates how individual postures are encoded in PCs 2, 4, and 5

To further explore what types of differences between individual flies might be encoded in the scores for PCs 2, 4, and 5, we systematically searched for an axis in this PC subspace that resulted in symmetric changes in posture with regard to the longitudinal axis of the body. We found an axis which describes the mean distance of the leg tips to the fly body (Fig. 7B). The contribution of PC 4 is very weak, which is not surprising since we searched for highest symmetry and PC 4 describes asymmetric covariations. Interestingly, flies 1, 2, and 3, which we already referred to in Figures 2 and 5, are all close to the symmetry axis and the qualitative differences in their postures, which mainly involved how sprawled these flies walked, are represented well in the artificially produced postures shown in Figure 7A. PC 4 on the other hand seems to describe less symmetric differences between individuals as presented for Fly 4 in Figure 5. However, this example demonstrates that PCs 2, 4, and 5 can be used to quantitatively differentiate between the idiosyncrasies of individual flies, potentially enabling grouping of similar flies to enhance signal to noise ratios or even to compensate for effects measured in multiple individuals by computationally subtracting their idiosyncrasies.

## Discussion

In this study, we used an extensive data set of a large number of flies and many instances of walking sequences to investigate the naturally occurring intra- and inter-individual variability of walking behavior on the level of leg tip kinematics. Initial qualitative observations in this data set readily replicated anecdotal impressions from previous studies that flies walk in an idiosyncratic manner (Fig. 2). A quantitative analysis of leg kinematics using principal components analysis (PCA) revealed that indeed much of the observed behavioral variability can be captured by lower-dimensional representation of five PCs (Fig. 3A). In this regard, two principal components (PCs 1 and 3) captured general variability aspects across individuals related to invariant movement patterns, such as the general positional changes during a step cycle, and interleg coordination (Fig. 3B and D, Fig. 4B and D); at the same time, three other PCs (2, 4, and 5) were related to and described individual-specific aspects of walking behavior (Fig. 3C, E, and F, Fig. 4C, E, and F). Taken together, the PCA approach used here was able to separate these two different sources of variability into distinct sub-domains (Fig. 5). In this notion, especially the contributions of the individual-specific PCs can be regarded as a compact fingerprint of a fly’s idiosyncratic way of walking (Fig. 5E) that is distinct from more general features shared by all flies. Finally, a further exploration of this individual-specific PC space revealed that general high-level features, like kinematic changes after experimental interventions (Fig. 6) or the overall postural width (Fig. 7) can be compactly detected, described, and quantified.

Postural variability in the present data set occurs on three different levels – the intra-step level, the step-to-step level, and the inter-individual level. The intra-step level thereby refers to positional variations of the tarsal tips over time; its main component is essentially the periodic back and forth movement of all legs during swing (protraction) and stance (retraction) movements. The specifics of these movements are mainly captured by PCs 1 and 3, with PC 1 reflecting the canonical tripod coordination and PC 3 reflecting deviations from this tripod coordination. Because this dynamical component of variability is large as compared to more subtle aspects of walking, like overall posture or left-right asymmetries, it is not surprising that PC 1 is the most prominent PC and generally captures a large part of the walking pattern across individuals. Together with PC 3 it therefore is suited to compactly represent the overall coordination an animal is using. So far, there have been only few attempts to efficiently characterize coordination of the six ambulatory legs of walking insects in a more compact form; TCS is one measure (Wahl et al., 2015; Wosnitza et al., 2012), but it has limitations, especially if the walking pattern deviates strongly from tripod coordination. A characterization based on these two PCs, whose contributions also seem to be negatively correlated with each other (see Fig. 4B and D), could serve as an alternative approach to efficiently describe coordination in future studies. An aspect that was not yet analyzed in detail here is the temporal variation of the contribution of these two PCs, their score time courses. Since PC 1 and 3 captured the actual movement of the legs, the scores of these two PCs will be modulated periodically over the time course of individual legs. Preliminary analysis of these scores indeed revealed this to be the case, but this was not explored in detail yet. Further examination in this regard could reveal more subtle relationships between how PC 1 and 3 are modulated in the context of interleg coordination and even how individual flies combine these two motor patterns that establish a basic tripod coordination (PC 1) and deviations from it (PC 3).

In contrast to PCs 1 and 3, PCs 2, 4, and 5 capture more static and inter-individual postural differences, as indicated by the results shown in Figures 4, 5 and 7. It is evident that the postures of individual flies occupy different parts of this PC subspace. Consequently, this subspace should be helpful to describe these individual differences in the first place, but its usefulness can be readily expanded. We explored two exemplary expansions here to some extent. The first was an experimental intervention that introduced known changes in leg kinematics which, in turn, were picked up clearly in the space spanned by PCs 2, 4 and 5. The second expansion was a top-down search in this space that describes overall changes in postural width. This approach can be useful in other novel interventions, in which putative effects are to be detected but for which a more unbiased analysis is desirable or in which several effects combine in a subtler way. Previous studies focused on obvious kinematic parameters like stance amplitude, durations of swing or stance phases, or AEP and PEP positioning. While these singular measures are certainly informative, a more unbiased way of looking at putative effects of interventions might reveal other effects that are less intuitive, more complicated, or interdependent.

In this context it has to be mentioned that PCA has the general limitation of only being capable of capturing linear correlations between the analyzed variables. It is conceivable that more complicated approaches for dimensionality reduction will yield more concise, compact, or clearer description and separation of the different levels of variability contained in the present data set. A previous study, for instance, used Uniform Manifold Approximation and Projection (UMAP) to explore the high-dimensional kinematic data of walking flies and found similar reductions of these data (DeAngelis et al., 2019). However, the analysis we carry out here already captures many interesting and helpful aspects of variability and the strength of PCA in the present context is its intuitive interpretability with regard to what individual PCs mean for posture and movement.

Another important point about our study is that we focused on straight walking in a relatively narrow speed range and restricted our analysis to isogenic male flies of the same age and reared in identical conditions. These restrictions were intentional, as we wanted to first establish the general approach and its usefulness for a more controlled subset of all possible behavioral data. Furthermore, we wanted to exclude additional sources of potential variability, based on parameters like walking speed, age, or sex, to name a few. However, even in this relatively controlled data set we nevertheless found a quite obvious diversity of idiosyncratic ways of walking in the flies investigated here. It is nevertheless true, that our analysis necessarily only reflected kinematic details which were contained in the selected data. Natural extensions of this analysis might focus on systematic influences of walking speed, curve walking, or age, among others. Expanding the dimensionality step by step and in a controlled way should successively elucidate the full spectrum of natural variability in fly walking behavior, thereby enabling more precise investigation of the fundamental principles underlying the motor control of insect walking.

One aspect which would be manifest to next focus on is, of course, walking speed, since it indeed has strong effects on many kinematic parameters and interleg coordination in walking insects (DeAngelis et al., 2019; Strauß and Heisenberg, 1990; Szczecinski et al., 2018; Wosnitza et al., 2012). Here, we chose to intentionally exclude speed to be able to more closely focus on individual differences between flies as well as general invariant dynamics of straight walking, as captured by PCs 1 and 3. However, one interesting connection between walking speed and variability that was anecdotally observed in previous studies and that we also qualitatively noticed in the present data set (data not shown), is that AEP and PEP positioning as well as phase relationships between legs on average become stricter, i.e. less variable, the faster a fly walks. This has not yet been quantified systematically, nor is there a strong hypothesis what the cause for this is. While we assume that this is a general effect, there might also be an individual component to this speed dependence of variability which, in turn, might hint at inter-individual differences in the neural control of walking. The approach presented here should be readily expandable to investigate this issue. Even more pronounced and qualitative shifts in the specific makeup of PCs and their contributions are expected when not only straight walking is accepted into the data set, but also curve walking. In insects, curve walking entails fairly strong changes in the kinematics of all legs as indicated by previous studies, mainly on stick insects (Du□rr and Ebeling, 2005; Gruhn et al., 2008). All legs contribute in a specific kinematic manner to curve walking. These postural changes will be reflected in the detected PCs, adding yet another layer of variability that reflects invariant aspects of curve walking, on the one hand, and potential inter-individual differences in the way single flies implement this change in motor output.

Taken together, in this study we provide the first extensive study of variability in *Drosophila* walking behavior on the level of isogenic individuals. The combination of dimensionality reduction using PCA with the interleg coordination measure TCS and optogenetic inhibition experiments shed some light on the natural variability that is present in walking behavior and proposed an enhanced way of describing, comparing, and quantifying data of freely walking flies. We thereby hope to build the fundament for further systematic investigation of the principles underlying the natural variability, as we are convinced that this path will inevitably lead to a deeper and more complete understanding of how insect nervous systems produce highly functional, versatile, and robust walking behavior.

## Data availability statement

Data and code are available on inquiry to the corresponding author.

## Author contributions

V.G. carried out experiments. V.G. analyzed data and created figures. V.G., T.B., and A.B. conceptualized the study. V.G., T.B., and A.B. wrote the manuscript. A.B. provided funding.

## Acknowledgements

We would like to thank Michael Dübbert and Mehrdad Ghanbari of the electronics workshop for excellent technical support. We would like to thank Sima Seyed-Nejadi, Sherylane Seeliger, and Corinna Rosch for fly husbandry.

